# Formation and synaptic control of active transient working memory representations

**DOI:** 10.1101/2020.08.30.273995

**Authors:** Sophia Becker, Andreas Nold, Tatjana Tchumatchenko

## Abstract

Neural representations of working memory maintain information temporarily and make it accessible for processing. This is most feasible in active, spiking representations. State-of-the-art modeling frameworks, however, reproduce working memory representations that are either transient but non-active or active but non-transient. Here, we analyze a biologically motivated working memory model which shows that synaptic short-term plasticity and noise emerging from spiking networks can jointly produce a working memory representation that is both active and transient. We investigate the effect of a synaptic signaling mechanism whose dysregulation is related to schizophrenia and show how it controls transient working memory duration through presynaptic, astrocytic and postsynaptic elements. Our findings shed light on the computational capabilities of healthy working memory function and offer a possible mechanistic explanation for how molecular alterations observed in psychiatric diseases such as schizophrenia can lead to working memory impairments.

## Introduction

Working memory (WM) is indispensable for a variety of cognitive functions ranging from sensory processing and integration of information to behavioural planning and execution (Baddeley, 2012). The diverse role of WM and its interaction with other cognitive processes is reflected in the complexity of its neural representation. Traditionally, WM is thought to be represented by elevated, asynchronous spiking or regular, synchronized population spikes of the memory-encoding neuronal population during the delay period after removal of the sensory cue (‘persistent delay activity’) (Fuster and Alexander, 1971; Funahashi et al., 1989; Christophel et al., 2017; Barbosa et al., 2020; Mongillo et al., 2008; Murray et al., 2017). This view has been extended by three other types of neural WM correlates in the brain. The first is referred to as ‘sequential delay activity’, where the neurons spike asynchronously but in regular sequences across the delay period (Harvey et al., 2012; Schmitt et al., 2017; Druckmann and Chklovskii, 2012). Alternatively, items can be held in WM through short bursts with high firing that are coupled to global brain rhythms (‘bursting delay activity’) (Lundqvist et al., 2016, 2018; Bastos et al., 2018) or based on Hebbian short-term plasticity (Fiebig and Lansner, 2017). Finally, WM items can be encoded as a temporary change of synaptic properties that slowly relax back to their baseline during the delay period (synaptic traces) (Mongillo et al., 2008; Stokes, 2015; Masse et al., 2019; Barbosa et al., 2020). Since these synaptic traces do not encode WM as spiking activity, this representation is often referred to as ‘silent’ WM. In contrast to spiking representations, silent WM is ‘inactive’ (Olivers et al., 2011; Trübutschek et al., 2017) and not directly accessible for active processing (Trübutschek et al., 2019). Recent computational modeling frameworks have reconciled experimental evidence for both persistent and silent WM representations (Mongillo et al., 2008; Trübutschek et al., 2019; Masse et al., 2019) and sequential and persistent WM (Druckmann and Chklovskii, 2012; Orhan and Ma, 2019).

These models produce either infinitely long, spiking memory representations or finite but silent WM representations. For example, the persistent, sequential and bursting delay activity WM models maintain the WM infinitely. The memory can only be terminated by an external off-signal, for example a population-specific inhibitory input (Barbosa et al., 2020) or a global reduction of excitatory input (Mongillo et al., 2008). Biologically, these off-signals could correspond to a loss of attentional focus on the WM representation (LaRocque et al., 2014), a local regulatory signal from inhibitory circuits (Lewis et al., 2004; Shilyansky et al., 2010) or a top-down input from other brain regions (Edin et al., 2009). But experiments show that WM activity is always characterised by its finite duration (Jonides et al., 2008), a feature that distinguishes it from intermediate and long-term forms of memory. This indicates that the maintenance period of working memory activity has a natural upper limit. Current models of spiking WM do not account for finite, spiking WM representations. In contrast to active representations, the synaptic traces of silent WM models (Mongillo et al., 2008) naturally decay over several hundred milliseconds to seconds and are therefore inherently transient. However, they do not allow to update the information held in WM to the same degree as models of spiking WM representations (Trübutschek et al., 2019). This raises the question whether we can build models that produce an actively spiking, yet intrinsically transient WM representation.

Studying pathologies with WM maintenance deficits provide insights into how such models can be built mechanistically. For instance, ablating the presynaptic molecule CHL1, which supports vesicle recycling in synaptic transmission, reduces the duration over which mice can maintain WM (Kolata et al., 2008). Through its influence on presynaptic vesicle recycling, CHL1 modifies the short-term plasticity (STP) dynamics at the synapse. Synaptic STP regulates synaptic transmission efficacy across several milliseconds to minutes (Zucker and Regehr, 2002) and arises from the interaction of presynaptic vesicle and calcium dynamics (Zucker and Regehr, 2002; Tsodyks et al., 1998). Recordings in the rat medial prefrontal cortex show firing rate-dependent synaptic facilitation and depression during WM (Fujisawa et al., 2008). These findings confirm computational predictions about the role of synaptic STP in silent WM (Mongillo et al., 2008; Masse et al., 2019). Similar to CHL1 signaling, other molecular signaling cascades also affect STP at the synapse (Zucker and Regehr, 2002; Moresco et al., 2003; Lovinger, 2008; Von Engelhardt et al., 2010; De Pittà et al., 2016) and can, if they are impaired, disrupt synaptic transmission and WM dynamics. Such malfunctions of WM are at the heart of several psychiatric diseases, including schizophrenia (Frydecka et al., 2014; Fioravanti et al., 2012), autism (Kercood et al., 2014), depression (Christopher and MacDonald, 2005; Rose and Ebmeier, 2006) and ADHD (Kasper et al., 2012; Matt Alderson et al., 2013). For example, a hallmark of schizophrenia are abnormal glutamatergic (Coyle, 2006) and GABAergic (Belforte et al., 2010) neurotransmission as well as altered dopaminergic signaling (Van Snellenberg et al., 2016). Since WM function strongly depends on these messenger cascades (Van Snellenberg et al., 2016), molecular alterations are believed to play an important role in compromising WM in schizophrenia Harrison and Weinberger (2005); Coyle (2006). Specifically, these lead to disinhibition in neurons and synapses and, thereby, cause an excitation-inhibition imbalance on the network level that interferes with healthy working memory function and cognitive processing in general (Yizhar et al., 2011). Many of these molecular alterations that have been associated with WM impairments and psychiatric disorders more generally are of genetic origin (Harrison and Weinberger, 2005). One putative genetic link between disrupted synaptic signaling and WM dysfunction is the single-nucleotide polymorphism R345T/mutPRG-1 (Vogt et al., 2016) that occurs in roughly 0.6 percent of the European and US population. It affects the plasticity-related gene 1 (*prg1*) and leads to dysfunction of the PRG1 protein and the associated signaling cascade.

The PRG1 protein is expressed at the postsynaptic density of glutamatergic synapses (Bräuer et al., 2003) in several brain regions, including the hippocampus and the somatosensory cortex (Trimbuch et al., 2009; Unichenko et al., 2016). Together with the astrocytic enzyme autotaxin (ATX), PRG1 regulates the presynaptic release probability via transmission-dependent changes in the concentration of the lipid messenger molecule LPA in the synaptic cleft (Yung et al., 2015). In the following, we refer to this mechanism as PRG1 mechanism. Humans with PRG1 impairments show altered sensory integration (Vogt et al., 2016). In mice with an equivalent of the human mutation of PRG1, the dysfunction of the PRG1 mechanism produces pathologically high presynaptic release probabilities (Trimbuch et al., 2009) and impaired short-term synaptic plasticity in glutamatergic synapses (Unichenko et al., 2016). At the network level, this produces an overall shift of the excitation-inhibition balance towards excitation, facilitating hyperexcitability (Trimbuch et al., 2009) and changes in both the spontaneous and the evoked activity of neuronal networks (Unichenko et al., 2016; Vogt et al., 2016). The altered cellular and network function due to PRG1 impairments manifests in the behaviour of PRG1-KO mice as frequent epileptic phenotype (Trimbuch et al., 2009), reduced resilience to stress (Vogt et al., 2016) and – as in humans with PRG1 dysfunction – altered sensorimotor processing (Unichenko et al., 2016; Thalman et al., 2018). These conditions link PRG1 impairments to schizophrenia (Yizhar et al., 2011), such that PRG1-KO mice are increasingly used as an animal model for schizophrenia and other psychiatric disorders with similar symptoms (Vogt et al., 2016; Thalman et al., 2018). Computational models allow us to explore the impact of synaptic signaling on WM dynamics and can inform experimental research investigating WM function in health and disease.

The impact of the PRG1 mechanism on working memory dynamics can be studied in a computational model that integrates synaptic short-term plasticity and spiking network dynamics (Mongillo et al., 2008). The defining element of this model is a network nonlinearity introduced by synaptic short-term plasticity (Tsodyks et al., 1998) that allows two WM states to coexist (bistability) (Mongillo et al., 2008). The initial state of the network determines whether its dynamics settle into a stable limit cycle or a stable fixed point. The former is characterised by a regularly oscillating firing rate, the latter by asynchronous low-rate firing of the memory population. This bistability can be leveraged as a WM model since a short, sufficiently strong excitatory input into a small subpopulation (the WM ‘cue’) can switch the network dynamics from its stable low-rate state into the oscillatory regime, that is interpreted as an active spiking WM of the input. We build on this WM model and integrate synaptic PRG1 signaling.

In our work, we show that short-term synaptic plasticity allows for active transient delay activity during a working memory protocol in addition to silent and persistent active WM regimes. PRG1 allows for a modulation of the duration of the transient active representation via astrocytic and postsynaptic signaling. We identify intrinsic noise as a key mechanism that maintains or disrupts the active working memory representation in a firing rate model. The duration of transient active WM is robust to changes in network size, but sensitive to network connectivity. It constitutes a natural upper limit for WM maintenance that could be altered in psychiatric disorders.

## Results

### A synaptic working memory model with postsynaptic and astrocytic signaling

The interaction of the postsynaptic protein PRG1, the astrocytic enzyme ATX and the lipid signaling molecule lysophosphatidic acid (LPA) in the synaptic cleft modulates presynaptic short-term plasticity (STP) in glutamatergic excitatory synapses (Trimbuch et al., 2009; Unichenko et al., 2016). To model the influence of this PRG1 mechanism on synaptic STP, we build on the STP synapse model by Tsodyks et al. (1998). Synaptic STP comprises temporary changes in synaptic efficacy that arise from the competing effects of fluctuating neurotransmitter availability and calcium binding at the presynaptic release sites (Zucker and Regehr, 2002). Upon arrival of a presynaptic spike, calcium binds at the active zones and leads to release of neurotransmitter vesicles. In between spikes, calcium slowly unbinds and the neurotransmitter storage is refilled. When the depletion of neurotransmitter due to incoming spikes exceeds the rate of replenishment, then this leads to a decrease of neurotransmitter release upon presynaptic spike arrival (short-term depression). This effect is counteracted by calcium binding that increases neurotransmitter release (short-term facilitation). These synaptic dynamics are captured by two equations (Figure 1 B, upper box), that describe the interaction of the presynaptic variables *x* and *y*: The relative amount of ready-to-release neurotransmitter vesicles *x* at the presynaptic active zones varies between 0 (complete depletion of vesicles) and 1 (full vesicle storage). Similarly, the relative amount *y* of calcium that is bound in the presynaptic active zones ranges between 0 (no calcium bound) and 1 (maximum amount of calcium bound). Neurotransmitter is replenished and calcium unbinds with time constants *τ*_*D*_ and *τ*_*F*_, respectively.

**Figure 1:**
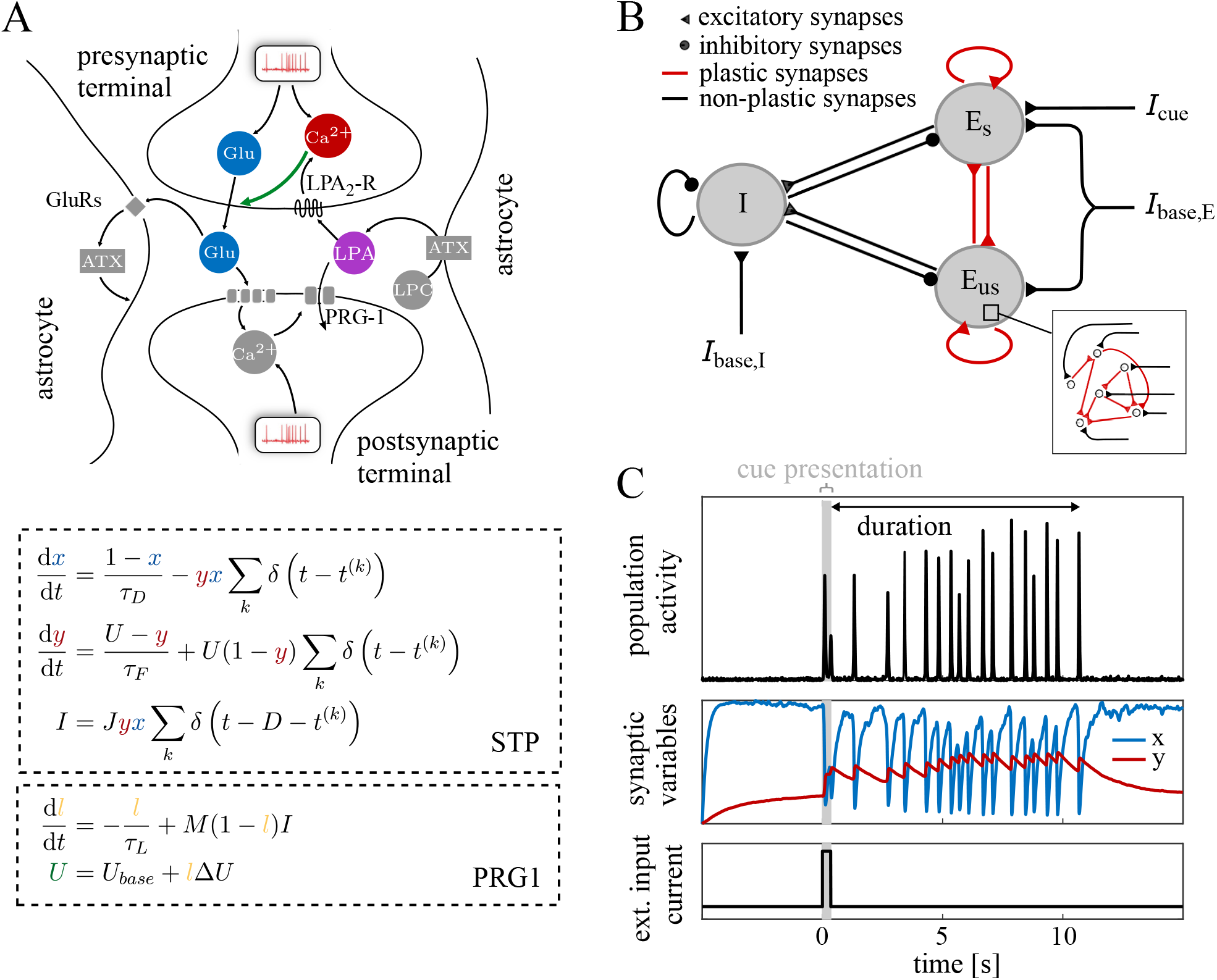
A synaptic working memory model with postsynaptic and astrocytic signaling. (A, top) Sketch of the transmission dynamics at glutamatergic synapses, as determined by the components of presynaptic STP and the transmission-dependent modulation by the astrocytic enzyme ATX and the postsynaptic protein PRG1. Presynaptic STP influences synaptic transmission via glutamate availability (*x*) and calcium binding (*y*) dynamics at the presynaptic active zones.The astrocytic and postsynaptic components of the PRG1 mechanism modulate presynaptic calcium binding rates via LPA binding (*l*). (A,bottom) shows the corresponding mathematical description of the synaptic dynamics, see text for details. The resulting synaptic current *I* is transmitted with a temporal delay *D*. (B) Architecture of the spiking network, comprising one inhibitory and two excitatory populations. One excitatory population is selective (E_*s*_) to the WM cue *I*_*cue*_, the other one (E_*us*_) is unselective. Black lines mark synapses with constant weights and red lines correspond to synapses equipped with STP dynamics, either with or without the PRG1 mechanism. All network populations receive an external baseline input current. (C) Transient WM dynamics, here simulated in an STP rate model. Upon the presentation of the cue, the memory population maintains the WM actively as synchronized population activity before it transitions into a silent representation with slowly decaying synaptic WM traces in *x* and *y*. The duration of the transient population activity corresponds to the active WM duration.

As depicted in Figure 1 A, the PRG1 mechanism interferes with STP dynamics by modulating the rate at which presynaptic calcium binds upon spike arrival. This effect is mediated by temporary increases in the concentration of LPA in the synaptic cleft that allow LPA to bind to its presynaptic LPA2 receptors (Trimbuch et al., 2009). Upon LPA binding, the G-protein coupled LPA2 receptors produce intracellular calcium responses (Vogt et al., 2017; Berridge et al., 2003). More calcium then binds to presynaptic active zones and, in consequence, more glutamate is released in response to a presynaptic spike. The LPA concentration in the synaptic cleft is modulated by postsynaptic and astrocytic signaling in a transmission-dependent way. When a spike is transmitted, the glutamate released into the synaptic cleft binds to receptors in the astrocyte (Thalman et al., 2018) and the postsynaptic terminal (Yung et al., 2015). The former boosts the synthesis of new LPA molecules by increasing the activity of the astrocytic enzyme ATX which synthesizes LPA from its precursor lysophosphatidyl choline (LPC) (Keune et al., 2016). The glutamate that binds to postsynaptic receptors inhibits the activity of the postsynaptic trans-membrane protein PRG1 via calmodulin (CaM) signaling (Tokumitsu et al., 2010). PRG1 usually takes up LPA from the cleft into the postsynaptic cell (Trimbuch et al., 2009). Due to the inhibition of PRG1 activity upon CaM binding, the new LPA molecules can remain in the cleft and bind to LPA2 receptors, where they affect presynaptic calcium dynamic as described above. The interplay of the astrocytic synthesis of LPA, the inhibition of PRG1-mediated postsynaptic LPA uptake and the effects of LPA binding on presynaptic calcium dynamics constitute the transmission-dependent PRG1 mechanism. In contrast to the effects of classic synaptic STP, which affects synaptic dynamics in the millisecond- to second-range (Zucker and Regehr, 2002), the LPA-mediated effects in the presynapse introduce longer timescales (Berridge et al., 2003). We integrate the PRG1 mechanism into the STP synapse model (Tsodyks et al., 1998) by adding a variable *l* that represents the amount of LPA bound by the presynaptic LPA2 receptors (Figure 1 B). It is driven by the amount of LPA available in the synaptic cleft, which is subject to transmission-dependent postsynaptic and astrocytic modulation. Therefore, the increase in *l* depends linearly on the transmitted synaptic current *I* and the strength *M* of postsynaptic and astrocytic modulation. The gradual unbinding of LPA from the LPA2 receptors is expressed as a decay of *l* with time constant *τ*_*L*_. Bound LPA *l* increases the presynaptic calcium binding rate *U* by *l*Δ*U*. This reflects the modulation of presynaptic STP dynamics and the associated release rate of the synapse by the PRG1 mechanism.

To study the impact of STP and PRG1 synaptic dynamics on WM activity, we consider spiking networks that comprise two excitatory and one inhibitory population. Each population consists of spiking leaky integrate-and fire (LIF) neurons that are randomly and sparsely connected. The evolution of the membrane potential of a neuron *i* over time is therefore given as

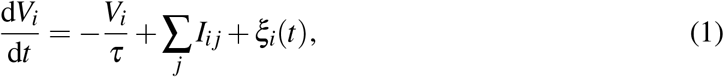

where *τ* is the membrane time constant, *I*_*i j*_ is the current transmitted through the synapse from neuron *j* to neuron *i* and *ξ*_*i*_ is an external white noise input (see Methods). When *V* reaches a threshold value *V*_*θ*_, the neuron emits a spike and the potential is reset to its resting potential. In addition to the white noise baseline input, a short ‘cue’ input current is presented to the smaller of the two excitatory populations. This population represents the memory-encoding, ‘selective’ population. It is embedded into the surrounding E-I network that comprises the larger, ‘unselective’ excitatory population and the global inhibitory population (Figure 1 C). We consider two types of spiking networks which differ in their E-to-E synapses: one includes Tsodyks-Markram STP synapses (Tsodyks et al., 1998) (STP networks, see Mongillo et al. (2008)) and the other includes PRG1-modulated STP synapses as described above (PRG1 networks). The synaptic efficacies between populations and the remaining network parameters are chosen such that excitatory and inhibitory inputs into each population are balanced (global E-I balance (Van Vreeswijk and Sompolinsky, 1998)). We explore the underlying mechanisms of the spiking network behavior in a STP firing rate model. In contrast to the spiking networks, the rate model consists of only one excitatory population that receives external baseline input as well as the WM cue. The external inputs have negative values to mirror the inhibitory inputs received by excitatory populations in the spiking networks (see Methods). Figure 1 D shows an example of the transient WM dynamics arising in STP and PRG1 networks, depicted for an STP rate network. The WM is maintained actively via repeated synchronized spiking of the memory population before transitioning into a silent representation, where the memory is contained in a decaying synaptic trace.

### Transient WM activity duration is modulated by synaptic dynamics

STP and PRG1 network simulations reveal five dynamical regimes (Figure 2 A). After the presentation of the WM cue, the excitatory selective population either (i) returns to asynchronous firing of the same rate as before the cue (‘silent’, see e.g. Mongillo et al. (2008)), (ii) shows a finite number of synchronized population spikes before returning to pre-cue asynchronous firing (‘transient’), (iii) enters sustained synchronized population spiking (‘persistent’, see e.g. Mongillo et al. (2008)), (iv) exhibits a chaotic regime with fluctuating levels of synchronization (‘self-evoked’, see e.g. Cortes et al. (2013)), or (v) produces asynchronous firing at rates that are elevated above the low-rate baseline (‘asynchronous’, see e.g. Brunel (2000)). As depicted in Figure 2 A, we traverse through these five regimes by increasing the baseline input current. For the remainder of our study, we focus on the transient WM regime and how its dynamics are affected by changes in synaptic parameters.

**Figure 2:**
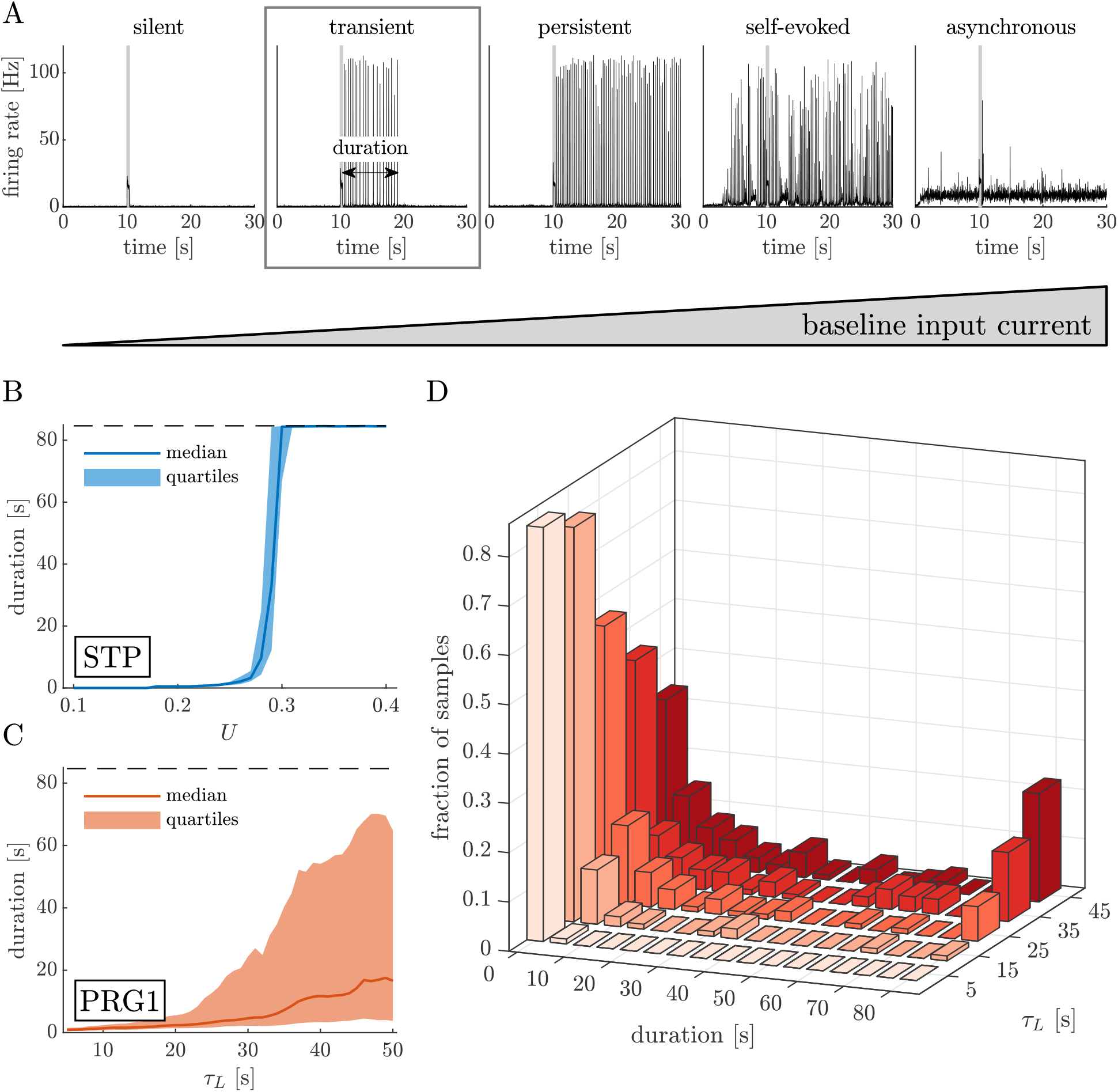
Transient WM activity duration is modulated by synaptic parameters. (A) Neural networks with short-term plastic STP and PRG1 synapses transition through five dynamical regimes as the baseline current into the network increases. For low baseline inputs, the memory population maintains the WM of the cue as a silent synaptic trace (left, Supplement, see Mongillo et al. (2008)). As the baseline current increases, the memory population exhibits first transient (2nd from the left), then persistent (center) population spike activity in response to the WM cue. While the transient WM regime automatically transitions to a silent synaptic representation (Supplement), the persistent regime can only be terminated by an off-cue (Mongillo et al., 2008). Higher external baseline inputs lead to self-evoked activity (2nd from the right, see Cortes et al. (2013)) and, finally, to elevated asynchronous activity (right, see Refs. (Mongillo et al., 2008; Brunel, 2000)) that arises even before the cue presentation. In this article, we focus on the transient spiking WM state. (B) In networks with synaptic STP, the fraction of presynaptically recruited calcium per spike (*U*) controls whether the WM cue is represented by a silent synaptic trace, transient or persistent population spiking. Within a small window of values, *U* modulates the median duration of the transient, actively spiking WM representation. (C) The postsynaptic and astrocytic feedback of the PRG1 mechanism also regulates the duration of transient active WM representations. (D) Distribution of transient WM durations as a function of *τ*_*L*_. The *τ*_*L*_-mediated increase of the median transient WM duration in (C) is driven by the tail of the duration distribution, i.e. a more frequent occurrence of longer transient WM durations.

In the transient WM regime, the cue evokes periodic, synchronized firing in the excitatory selective population. After a finite period, the activity spontaneously transitions back to asynchronous low-rate firing and the memory population maintains the information as a decaying synaptic trace. No external off-cue is required to terminate the transient WM activity. A weak reminder stimulus can then ‘activate’ the memory and produce a population spike, similar to the silent WM regime (Figure S1). In contrast to the self-evoked regime, which can also exhibit spontaneous periods of transient population spike activity, the transient activity regime is characterized by regular spiking and the fact that transient population spikes need to be evoked by a cue.

Analogous to increases in baseline input current, changes in synaptic dynamics also induce a transition from the silent through the transient to the persistent WM regime. In this transition, the duration of the active WM representation continuously increases from zero to infinity. In STP networks, the transition is produced by varying the presynaptic calcium binding rate *U* (Figure 2 B) or the time constants of calcium unbinding *τ*_*F*_ and neurotransmitter refill *τ*_*D*_ (Figure S2 A, B). In the PRG1 network, the duration of transient WM activity is modulated by changes in the time constant of LPA unbinding *τ*_*L*_ (Figure 2 C). Similarly, the duration of the transient representation also increases with the effective LPA-binding rate *M* and the LPA-mediated change in presynaptic calcium binding Δ*U* (Figure S2 C, D) in the PRG1 network. Postsynaptic and astrocytic feedback of the PRG1 mechanism modulates transient WM duration via its influence on the presynaptic calcium binding rate *U* (Figure S3). The exact duration of the transient population spike activity is subject to trial-by-trial variability that increases with the median of transient durations (Figure 2 B). Interestingly, the increase in median duration is mainly driven by the tail of the distribution, i.e. by a higher incidence of long transient representations (Figure 2 D).

### Transient WM activity emerges from silent and persistent network dynamics due to noise

Which network mechanism leads to active transient WM representations? We approach this question in an STP rate network, which allows for a simplified and mathematically tractable analysis.

We find that an STP rate model without noise supports infinitely long population activity but does not exhibit transient cycles, which is consistent with previous reports (Mongillo et al., 2008). Apart from the lack of transient cycles, the noise-free STP rate network exhibits similar dynamical regimes as the spiking STP network. For weak baseline current, it converges to a stable steady-state with low population activity (*E*_*base*_ < −3.25, Figure 3 A) that allows for silent WM representations as decaying synaptic traces. For intermediate baseline inputs (−3.25 < *E*_*base*_ < −2.5, Figure 3 A), the initial conditions of the network determine whether its dynamics converge to the low-activity fixed point or a limit cycle around the upper, unstable branch of Figure 3 A. In this case, a sufficiently strong WM cue can switch the network dynamics from the low-activity fixed point into the limit cycle, which corresponds to the persistent WM regime in the spiking network. Higher current inputs lead to a stable limit cycle (−2.5 < *E*_*base*_ < −2, Figure 3 A) and, finally, to a stable, high-activity fixed point (*E*_*base*_ > −2, Figure 3 A). The latter mirrors the asynchronous regime in a spiking STP network.

**Figure 3:**
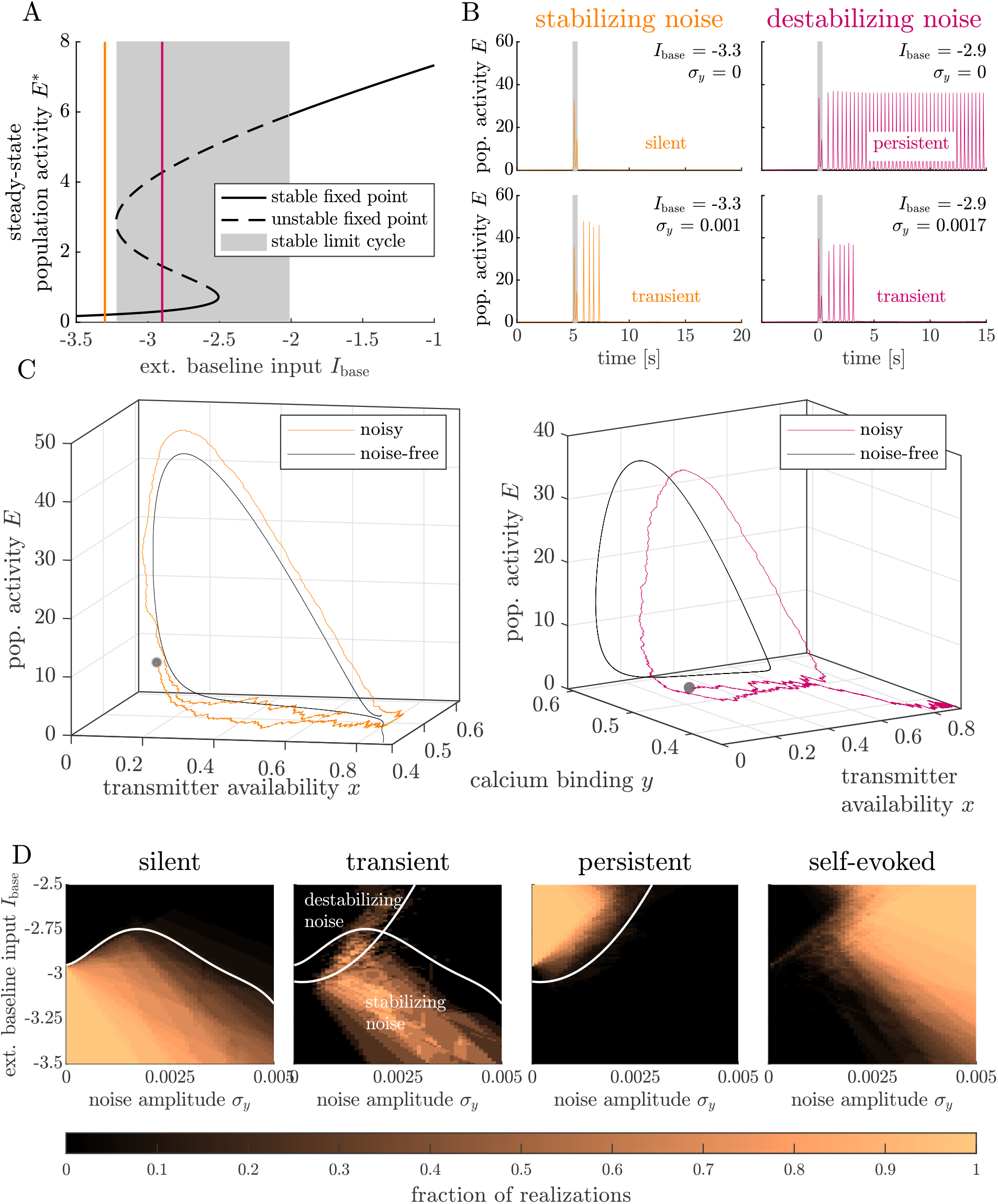
Transient WM activity emerges from silent and persistent network dynamics due to noise. (A) The phase space of the STP rate network exhibits a stable fixed point at low baseline currents. An STP network in this regime shows silent WM, corresponding to Figure 2 A (left) in the spiking network. Increasing baseline current leads the network dynamics through a bifurcation into a bistable regime (grey-shaded area), where the low-rate fixed point coexists with a persistent limit cycle. Presenting a sufficiently strong WM cue to networks in this regime evokes persistent WM activity (equivalent in the spiking network: Figure 2 A, middle). For even higher baseline currents, the bistability gives way to a new high-activity fixed point that is unaffected by the WM cue (Figure 2 A, right in the spiking STP network). In the silent regime prior to the first bifurcation, noise can stabilize a cue-evoked transient limit cycle (B, stabilizing noise). The presence of noise in the regime of persistent limit cycles can also lead to transient oscillations (B, destabilizing noise). (C) Typical trajectories of transient limit cycles emerging from stabilizing (orange line, left) and destabilizing noise effects (magenta line, right) at baseline input levels *I*_base_ = −3.3 (left) and *I*_base_ = −2.9 (right). Black lines show representative examples of noise-free trajectories. (D) Noise amplitude *σ*_*y*_ and baseline current *E*_*base*_ determine the prevalence of each network activity regime.

The absence of transient active WM representations in the noise-free STP rate network and their presence in spiking STP networks suggests that the transient regime is generated by noise in the network. Indeed, by adding a small perturbation to one of the network variables *x, y* or *E*, we obtain transient population spike activity during the delay period. The transient WM regime emerges in two ways (Figure 3 B): (1) noise transiently stabilizes population spike activity during the delay period, where a noise-free network receiving similarly low baseline inputs would have returned to its low-activity fixed point; or (2) noise destabilizes the persistent population spike activity that arises due the WM cue in a network receiving intermediate baseline inputs. In both cases, the network perturbation is not strong enough to induce transient population spikes without the prior presentation of the WM cue. To illustrate the dynamics of case (1), we consider a stereotypical noise-free trajectory originating from a high-*y*, high-*x* regime (see black line in Figure 3 C, left panel). It evokes one population spike before returning to the fixed point, passing close to its initial state. At this point, noise can pushed the system into a second cycle, inducing another population spike (orange line Figure 3 C, left panel). The reactivation of the cycle is non-deterministic, and the exact number of oscillations before returning to the fixed point is therefore stochastic. In case (2), network dynamics are governed by two stable attractors: a low-activity fixed point and a limit cycle (black line, right panel of Figure 3 C). First, the WM cue pushes the network dynamics from the low-activity fixed point into the limit cycle. Sufficiently strong noise can destabilize this limit cycle by pushing the network dynamics back into the basin of attraction of the stable fixed point (purple line, right panel of Figure 3 C). The destabilization takes place during the low-activity phase of the cycle, when the system is close to the boundary separating the regions where dynamics are drawn in the limit cycle and the fixed point, respectively.

The main requirement for the existence of transient cycles is an appropriate level of network noise. Without noise, an increase in baseline current *E*_*base*_ induces an abrupt transition between the silent and the persistent regime (Figure 3 D, panel 1 and 3). For sufficiently strong noise levels (*σ*_*y*_>0.001), transient cycles appear at the transition between silent and persistent regime (around *E*_*base*_ = −3, Figure 3 D, panel 2). This is because noise levels are strong enough to either push the system into an oscillation after the WM cue (stabilizing noise case, for low baseline currents *E*_*base*_) or out of the stable limit cycle towards the low-activity fixed point (destabilizing noise, for elevated baseline currents *E*_*base*_). While we observe both stabilizing and destabilizing noise effects, stabilizing noise covers a larger parameter range (Figure 3 D). Very strong noise eventually causes the self-evoked regime to supersede the previous combination of silent, transient and persistent regimes (Figure 3 D, panel 4). In conclusion, the transient WM activity in the rate model reproduces the (cue-evoked) transient WM regime in the spiking network and can be clearly distinguished from the self-evoked regime with spontaneous transient episodes (see e.g. Cortes et al. (2013)).

### The duration of transient active WM is robust to changes in network size but sensitive to network wiring

The transient network dynamics we observe could be caused by finite-size effects and, therefore, may disappear in large networks. We show that transient WM activity in spiking STP and PRG1 networks, however, is not restricted to small neuronal networks (Figure 4 A, B). We compare two spiking STP networks of biologically plausible sizes of 10 000 neurons (as used in Figure 2) and 60 000 neurons, respectively, and rescale their synaptic weights and external inputs with the square root of the number of neurons as in Van Vreeswijk and Sompolinsky (1998) (Methods). As we approach the balanced state limit of large networks, this rescaling reduces the effects of neuronal nonlinearities on network dynamics and allows us to study the network dynamics arising from synaptic nonlinearities. As in the 10 000-neuron STP network (Figure 2 B), the existence and duration of transient WM activity the 60 000-neuron STP network is controlled by changes in the synaptic parameter *U*, which describes the fraction of presynaptically recruited calcium per arriving spike (Figure 4 A). Transient WM activity and synaptic modulation of its duration is thus preserved in larger networks. However, the combination of baseline input currents *E*_*base*_ and presynaptic calcium binding rates *U* that gives rise to the transient WM regime differs across the two networks (Figure 4 A, B). Additionally, the duration of transient WM activity shows a higher variability in the larger network than in the smaller network.

**Figure 4:**
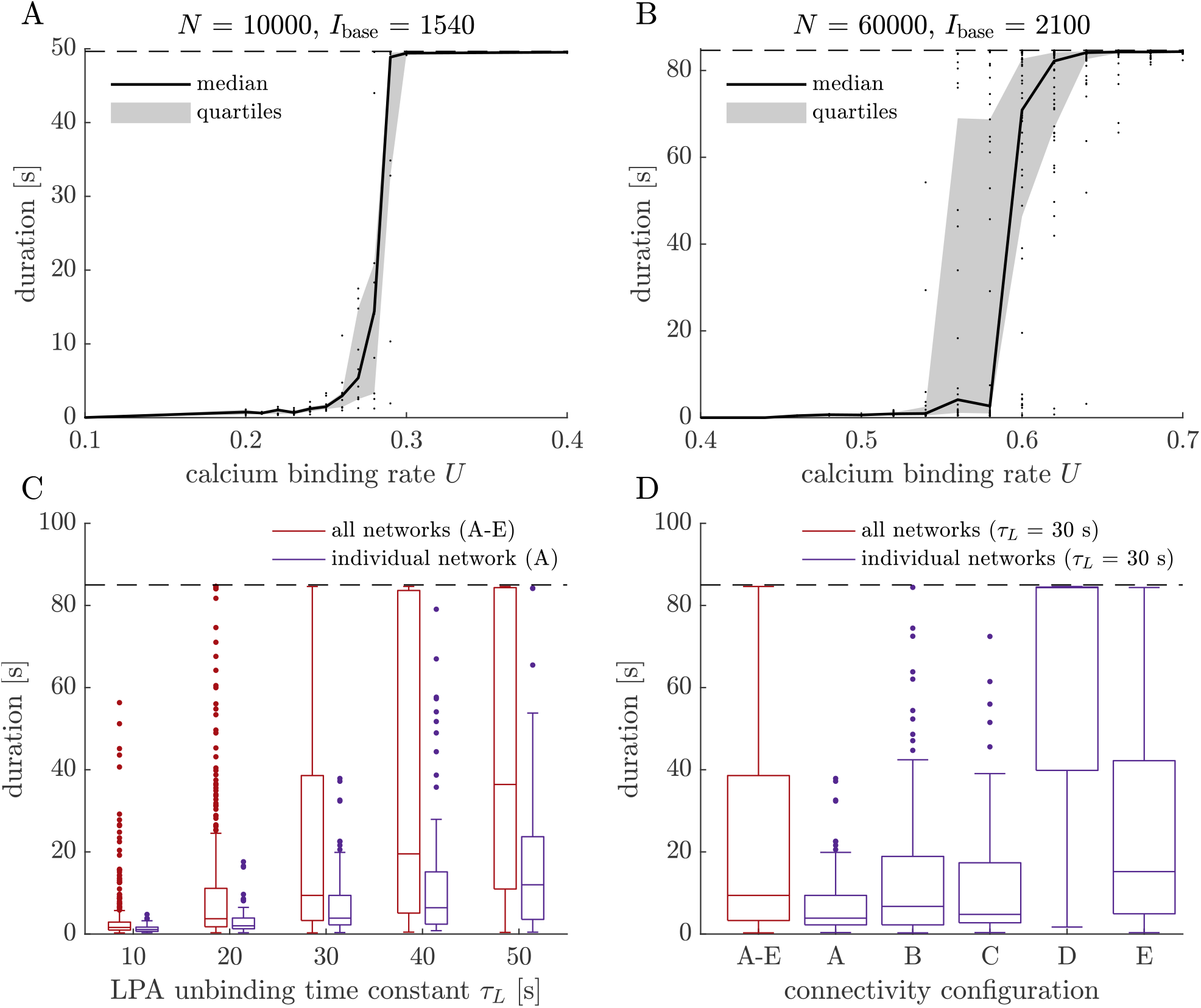
The duration of transient active WM is robust to changes in network size but sensitive to network wiring. (A, B) Networks with different sizes (number of neurons *N*=10 000 and *N*=60 000, consistent with cortical column sizes in different animals (Markram et al., 2015)) support transient WM. The transient regime appears in different ranges of the baseline input current *E*_*base*_ and the presynaptic calcium binding rate *U* in the two networks. (C, D) The wiring pattern of a network influences the statistics of the transient WM durations. For each of five networks with different wiring, 100 realizations are shown either separately for each network (purple boxplots) or together (red boxplot). Network wiring accounts for a large part of the variability of WM durations, but does not affect their trend to increase with the LPA unbinding time constant *τ*_*L*_ (C).

What is the source of this variability? So far, we have averaged the durations of transient activity across different instances of sparse random network connectivity. Could the resulting variations in network wiring have an influence on the duration or even the existence of transient WM activity in our STP and PRG1 networks? To this end, we compare the statistics of transient WM activity with and without variations in network wiring (Methods). The transient WM regime and the synaptic modulation of its duration is preserved across the networks we studied. Interestingly, differences in the wiring structure between the networks account for a large part of the variability in transient durations (Figure 4 C, D).

## Discussion

We identified a mechanism underlying transient spiking representations of working memory (WM) using a computational modeling framework. The existence of transient representations relies on synaptic short-term plasticity (STP) and its duration is regulated by STP as well as astrocytic and the postsynaptic feedback. We build upon the synaptic WM model by Mongillo et al. (2008) to include a post-to-presynaptic mechanism (Figure 1 A) based on the postsynaptic protein PRG1 (Trimbuch et al., 2009) and the astrocytic enzyme ATX (Vogt et al., 2016; Thalman et al., 2018). The effects of this signaling pathway introduce longer timescales (Berridge et al., 2003) to the fast dynamics of short-term synaptic plasticity (STP) (Zucker and Regehr, 2002). Integrating the PRG1 model synapses in a network model (Figure 1 B and C) allowed us to investigate the impact of the PRG1-modulated STP on WM dynamics.

The transient WM regime we report here consists of a finite period of population activity following the WM cue, and a subsequent representation as a silent synaptic trace (Figure 2 A). It shows that active and silent representations do not have to occur in a mutually exclusive fashion but can appear in the same memory trial. The transient activity regime allows processing during the first phase of active representation, and then switches to a energy-efficient silent mode. In contrast to previous models of active WM representations, such as presented by Mongillo et al. (2008), the transient WM regime does not require external stimuli to terminate the population activity. This is in line with the definition of WM (Baddeley, 2012) and offers a mechanistic explanation for naturally decaying working memory activity (Jonides et al., 2008; Pasternak and Greenlee, 2005). Electrophysiological studies in non-human primates typically use short delay periods (see e.g. Funahashi et al. (1989); Christophel et al. (2017)), and can therefore not be used to validate the upper limit of WM maintenance. The termination of WM activity during such experiments with short delay periods has been attributed to voluntary reorientation of attention to other WM items or distractors (Watanabe and Funahashi, 2014; LaRocque et al., 2014). The longer, self-terminated WM representation of the transient regime could therefore reflect an involuntary loss of attention over time.

The second key finding from our study is that the duration of transient WM activity is controlled by presynaptic, postsynaptic and astrocytic parameters of STP and the PRG1 mechanism (Figure 2 B-D). For example, a strong inhibition of postsynaptic PRG1 activity or a high production of new LPA by ATX (high parameter *M* in Figure 1 A) leads to high LPA binding (*l* close to 1) at the presynaptic side. This causes increased rates of presynaptic calcium binding upon spike and an increased synaptic release rate, which results in longer transient durations. In support of a synaptic modulation of WM maintenance, genetic ablation of the presynaptic molecule CHL1 in mice (Kolata et al., 2008) leads to reduced WM durations. CHL1 is involved in presynaptic vesicle recycling and therefore crucial for synaptic transmission. Like PRG1 impairments, CHL1 deficiency is associated with schizophrenia and altered cognitive ability, among others with respect to social behaviour and sensory-motor gating (Kolata et al., 2008). Sensitivity of WM duration to synaptic dynamics implies that synapses could be tuned to produce WM representations with different maintenance periods. This could allow the network to dynamically adjust WM durations to current computational needs. For instance, each astrocyte covers around 10^5^ synapses in mice and 2 *×* 10^6^ in humans (Allen and Eroglu, 2017), and controls synaptic transmission and plasticity (Sun et al., 2016; De Pittà et al., 2016). Based on our model, astrocytic signaling could therefore modulate the activity of a local ensemble of neurons that represent typical sub-features of WM items. This could support a feature-wise control of WM representations and their duration.

Which mechanism underlies the transient WM regime? We found that noise is the the main driver of transient WM dynamics in STP firing rate networks (Figure 3). Depending on the dynamical regime of the network, noise can take on two different roles. In a scenario where the noise-free network would return directly to low-rate baseline firing after the presentation of a cue (silent regime), sufficiently strong noise can produce noise-stabilized transient oscillations. Similar stabilizing effect of noise in networks have been reported in the context of statistical mechanics (Juel et al., 1997; Ushakov et al., 2005; Kiss et al., 2003; Morrone et al., 2011) and gene regulatory networks (Lee and Lee, 2018). In the neuroscience context, noise has been reported to stabilize rehearsal of long-term memories (Wei and Koulakov, 2014) and aperiodic attractors in networks of the olfactory system (Freeman et al., 1997). In the second scenario, where the cue stimulation is strong enough to switch the network dynamics into a stable limit cycle, noise destabilizes the persistent oscillation and causes firing rates to return to baseline population activity after a finite period. The idea that noise can destabilize memory representations is well known in computational neuroscience (see e.g. (Jonides et al., 2008)). Noise in our STP rate model is ‘internal’ network noise in the sense that it is generated by a stochastic perturbation of the synapse or network variables. Similar internal types of noise are known to influence spiking network dynamics. For example, we identified connectivity patterns as an important source of variability that influences the termination of population activity during WM delays.

In Figure 4 C and D, we show that the randomness of the network connectivity accounts for a large part of the variability of transient WM durations. This suggests that a plausible reason for differences in the WM representations across brain regions could be variations in their connectivity structure. But which features in the wiring structure prime networks to exhibit long or short transients? One possible approach to this question is to analyze the eigenvalue and rank structure of the connectivity matrix (Mastrogiuseppe and Ostojic, 2018; Schuessler et al., 2020). Alternatively, the prevalence of certain circuit motifs could be shaping network dynamics (Hu et al., 2018). Understanding the role of network wiring may open up the possibility to understand and leverage connectivity in complex network designs (Bouchacourt and Buschman, 2019). We also vary the network size and rescale the inputs and synaptic weights, such that large networks approach the balanced-state limit (Van Vreeswijk and Sompolinsky, 1998) and their dynamics are dominated by synaptic nonlinearities. We show that transient delay activity is preserved in such networks with 10^4^ and 6 *×* 10^4^ neurons (Figure 4 A and B). The transient WM regime is therefore likely caused by interactions between synaptic nonlinearities and internal noise.

Finite WM representations imply that forgetting is an integral part not only of long-term memory processes (Gravitz, 2019), but also of WM. Currently, two dominant theories of forgetting exist in the WM context. The first describes forgetting as a continuous decay of the WM representation over time, while the second attributes forgetting to the interference of the memory with other active perceptions or memories (Jonides et al., 2008). Here, destabilization through intrinsic network noise (Figure 3) indicates that internal noise sources corroborate the WM representation over time, eventually leading to WM loss. The sudden transition from active to silent WM representation is, at first glance, in conflict with EEG and fMRI recordings that show a gradually decaying activity signature in the memory-encoding brain areas (Pasternak and Greenlee, 2005). However, one WM item is represented by multiple, distributed subpopulations that encode separate features and subfeatures (Postle, 2006; Christophel et al., 2017) rather than by a single, isolated population. Then, many single memory populations that switch from active to silent representation after a stochastic period could combine to the gradual decay of the WM item. Rather than losing random neurons of its representation, the memory of the item becomes less and less detailed as more and more subfeatures are forgotten.

There are several ways in with our modeling work can be extended. First, the PRG1 synapse model can be adapted to include additional molecular pathways at the synapse, for example CHL1-related signaling. A first step towards this is to introduce separate variables for the post-synaptic and astrocytic elements of the PRG1 mechanism, that replace the shared parameter *M* (see Figure 1 A, Supplement). Future modeling work could also include neuronal and astrocytic morphologies as well as spatially structured connectivities to probe the behavior of synaptic ensembles. Experiments (Buschman et al., 2011) and modeling studies (Bouchacourt and Buschman, 2019) show that the WM performance decreases with the WM load, i.e. the amount of items held in WM. Studying transient WM in the presence of multiple stimuli could clarify whether this decrease is imposed by limitations in astrocytic or postsynaptic signaling. The methods introduced by Mi et al. (2017), which present an explanation for limited WM load in networks with classic short-term plasticity, could serve as a starting point.

To conclude, our modeling work presents a candidate mechanism for the finite working memory duration that is observed experimentally. Our findings emphasize the potential role of astrocytes in shaping transient working memory representations and points at mechanisms that may contribute to working memory malfunctions in psychiatric disorders such as schizophrenia.

## Acknowledgements

This study was supported by the Max Planck Society (T.T.), the German Research Foundation via CRC1080 (T.T.), the Joachim Herz Foundation (Add-on fellowship, A.N.), the German Academic Scholarship Foundation (S.B.) and the CRC 1080 Cooperation Program — “Molecular and Cellular Mechanisms of Neural Homeostasis” (S.B. and A.N.). We thank Johannes Vogt and Heiko Endle for valuable discussions and insights into the PRG1 signaling mechanism. We thank Sara Konrad for contributions to the neural network code and Pierre Ekelmans for stimulating discussions and constructive feedback.

## Author contributions

A.N. and T.T. conceived the study; S.B. and A.N. performed the research; S.B. prepared the first draft of the manuscript, S.B., A.N. and T.T. wrote and revised the manuscript.

## STAR Methods

### Spiking network model

In Table 1, we describe the spiking neuronal network model used in Figure 2 and Figure 4 according to the model reproducibility framework proposed by (Nordlie et al., 2009). The parameter values of the model are specified in Table 2. We implement the model in C++, post-processing and analysis of the simulation data are performed in Matlab 2020a (code available upon request).

**Table 1:** Spiking network model.

| Model summary |  |  |  |
| --- | --- | --- | --- |
| **Populations** | Three: excitatory unselective ($E_{us}$), excitatory selective ($E_s$), inhibitory (I) | | |
| **Connectivity** | Sparse, random connectivity |  |  |
| **Neuron model** | Leaky integrate-and-fire (LIF) neuron with fixed voltage threshold, reset potential and absolute refractory time |  |  |
| **Synapse model** | Two types in each network: non-plastic δ-synapse, short-term plastic synapse (either STP or PRG1) | | |
| **Input** | Independent white noise current input into each neuron, additional working memory cue input into neurons of $E_s$ | | |
| **Measurements** | Spiking activity in the excitatory selective population $E_s$, state variables of the plastic synapses connecting two excitatory selective neurons | | |

| Populations |  |  |  |
| --- | --- | --- | --- |
| Name | Elements | Size | Function |
| $E_{us}$ | LIF neurons | $N_{E,us}$ | Large excitatory population that is unselective to the WM cue |
| $E_s$ | LIF neurons | $N_{E,s}$ | Smaller excitatory population that is selective to the WM cue and can maintain a WM representation during the delay period |
| I | LIF neurons | $N_I$ | Global inhibitory population |

| Connectivity |  |
| --- | --- |
| **Randomness** | Random connectivity is generated by the following algorithm: For each potential connection between a target neuron $x_{i,\alpha}$ from population $\alpha$ and a source neuron $x_{j,\beta}$ from population $\beta$, we draw a random number from a uniform distribution between zero and one. If the number is below a threshold $p \in [0, 1]$, we establish a connection between the source and the target, otherwise the source and target remain unconnected. The total number of source neurons for each target neuron in a network of $N$ neurons therefore follows a binomial distribution with mean $p(N - 1)$. We call $p$ the connection probability of the network. |
| **Sparseness** | Neurons are sparsely connected in the following sense: The average number of incoming connections $C = p(N - 1)$ for each neuron is significantly smaller than the total number of neurons $N$ in the network, as $C/N = p \ll 1$. Due to the sparseness of the network connectivity, neurons receive inputs only from a relatively small number of neurons. The inputs into each neuron are therefore only very weakly correlated, such that we can neglect input correlations in sufficiently large networks. |
| **Self-/ Multi-connectivity** | Connections are only established between pairs of distinct neurons (no self-connectivity). Each pair of neurons can only be linked by a single synaptic connection (no multiconnectivity). |
| **Weights** | Fixed weights $J_{\beta,\alpha}^{(j,i)}$ for each connection between a source neuron $i$ in population $\alpha$ and a target neuron $j$ in population $\beta$. Connection weights are homogeneous within each synapse population, i.e. the set of synapses with common source and target neuron populations. The only exception are synapses emerging from $E_{us}$ into $E_s$: to reflect the variability of connection strength in very large synaptic populations like this one, each connection has probability $\gamma_{\text{pot}}$ to be assigned with a potentiated weight $J_{\text{pot}}$ instead of the baseline connection strength $J_{\text{base}}$. |
| **Delays** | All synapses connecting neurons within one population have non-zero synaptic delay times. For each such connection, a delay time is drawn uniformly at random from the interval $[D_{\min}, D_{\max}]$. |

| **Neuron model** |  |
| --- | --- |
| **Type** | Leaky integrate-and-fire (LIF) neuron. Each LIF neuron is characterized by its membrane potential $V(t)$. |
| **Subthreshold dynamics** | The dynamics of the membrane potential $V_i$ of an LIF neuron $i$ in population $\alpha$ between two consecutive spike times $t^{(k)}$ and $t^{(k+1)}$ are given by:<br>$\frac{dV_i}{dt}(t) = -\frac{V_i}{\tau}(t) + \sum_{j \neq i} I_{ij}(t) + I_{\text{ext},i}(t) \quad \text{if } t^{(k)} + t_{\text{ref}} < t < t^{(k+1)}$ $V_i(t) = V_r \quad \text{if } t^{(k)} < t < t^{(k)} + t_{\text{ref}}.$<br>Here, $\tau$ is the membrane time constant of the LIF neuron, $V_r$ is its reset potential and $t_{\text{ref}}$ is its refractory period. The neuron receives recurrent synaptic inputs $I_{ij}(t)$ through synapses from other neurons $j$ to neuron $i$ and an external white noise input $I_{\text{ext},i}(t)$. We assume that the membrane resistance $R$ is constant and can therefore be included in the representation of the currents. |
| **Spiking** | A neuron $i$ emits a spike when its membrane potential $V_i(t)$ crosses the threshold potential $V_\theta$. The time point of the $k$-th spike emission is saved as spike time $t^{(k)}$ and the membrane potential is set to the reset potential $V_r$. |

| Synapse models |  |
| --- | --- |
| Name | Dynamics |
| δ-synapse | A δ-synapse from neuron $j$ in population $\beta$ to neuron $i$ in population $\alpha$ transmits the synaptic current<br>$I_{ij}(t) = J_{\alpha\beta}^{(ij)} \sum_k \delta(t - D_{ij} - t^{(k)}),$<br>where $J_{\alpha\beta}^{(ij)}$ is the synaptic connection strength, $D_{ij}$ is the synaptic delay and $t^{(k)}$ is $k$-th spike time of the presynaptic neuron $j$. |
| STP synapse | See Figure 1 A, lower panel. |
| PRG1 synapse | See Figure 1 A, lower panel. |
| Input |  |
| **Baseline input** | Every neuron $i$ from population $\alpha$ receives an excitatory baseline input with constant mean $\mu_\alpha$,<br>$I_{ext,i} = \mu_\alpha + \xi_i(t),$<br>where $\xi_i(t)$ is Gaussian white-noise with mean zero and autocorrelation $\langle \xi(t)\xi(t') \rangle = \sigma_\alpha^2 \delta(t - t')$. |
| **Cue stimulation** | Neurons of the excitatory selective population $E_s$ receive a short, additional input current that represents a stimulation of the network with a working memory cue. After an initial period $T_{init}$ of baseline stimulation that allows the network to settle into stable dynamics, the mean and variance of the white-noise input into $E_s$ is increased by a factor $A_{cue}$ for a duration of $T_{cue}$. Afterwards, the white-noise input into $E_s$ returns to baseline stimulation for the remainder of the simulation. |
| Measurements |  |
| Type | Method |
| Population firing rate | Sum of spike counts in each population per second |
| Duration of population spike regime | Time between end of the cue stimulation and the last population spike. We use an in-built Matlab function for spike detection. |
| Statistics of population activity duration | Statistics of the duration of population spike activity are computed for a set of $S$ realizations of the same network. Sources of randomness that lead to differences across realizations are the random connectivity structure of the network, the initialization of synaptic and neuronal variables and temporal noise of the external input. The randomness in these sources is set via independent random seeds that are generated from a global random seed. The global random seed is chosen uniquely for each realization. |

**Table 2:** Standard parameters of the spiking network model. We deviate from these standard parameter values in the following cases. (1) The five dynamical regimes in Figure 2 A arise for different levels of baseline white-noise input, with means *µ*_*E*_ = *{*22.95, 23.165, 23.4, 23.85, 24.3*} mV/s* (subfigures Figure 2 A, from left to right). (2) We vary the synaptic STP parameters *U* = 0.1-0.4 in Figure 2 B, *τ*_*F*_ = 1-5 s in Figure S2 A and *τ*_*D*_ = 0-1.5 s in Figure S2 B. (3) Similarly, we vary the PRG1 synaptic parameters while keeping the STP parameters at their standard values as above: *τ*_*L*_ between 5 s and 50 s in Figure 2 C, D and Figure 4 C; *M* between 0 and 1 in Figure S2 C; and Δ*U* between 0 and 0.8 in Figure S2 D. (4) In Figure 4 A and B, we vary the synaptic STP parameter *U* as in case (2) for networks with two different sizes. In addition to the standard network above, we consider a larger network with *N* = 60 000, *N*_*E,us*_ = 43 200, *N*_*E,s*_ = 4 800 and *N*_*I*_ = 12 000. (5) For Figure 4 C and D, we compare the statistics of working memory activity durations with and without connectivity noise. To compute the statistics without connectivity noise, we fix the seed determining the random connectivity of each network realization at different values for network configurations A-E in Figure 4 D and only allow randomness in the external input and variable initializations.

| General simulation parameters | Values |  |
| --- | --- | --- |
| $dt$ – simulation time step | 0.1 ms | |
| recording bin size | 10 ms |  |
| $S$ – number of simulated realizations | 100 | |
| Populations parameters | Values |  |
| $N$ – total number of neurons | 10 000 | |
| $N_{E,us}$ – number of excitatory unselective neurons | 7 200 | |
| $N_{E,s}$ – number of excitatory selective neurons | 800 | |
| $N_I$ – number of inhibitory neurons | 2 000 | |
| Connectivity parameters | Values |  |
| $p$ – connection probability | 0.2 | |
| $J_{IE}$ – synaptic efficacy $E \rightarrow I$ | 0.135 mV/spike | |
| $J_{EI}$ – synaptic efficacy $I \rightarrow E$ | 0.25 mV/spike | |
| $J_{II}$ – synaptic efficacy $E \rightarrow I$ | 0.2 mV/spike | |
| $J_{base}$ – baseline level of $E \rightarrow E$ synaptic efficacy | 0.05 mV/spike | |
| $J_{pot}$ – potentiated level of $E \rightarrow E$ synaptic efficacy | 0.45 mV/spike | |
| $\gamma_{pot}$ – fraction of potentiated synapses | 0.1 | |
| $[D_{min}, D_{max}]$ – range of synaptic delays | [0.1 .. 1] ms | |
| Neuron parameters | E | I |
| $V_\theta$ – spike emission threshold | 20 mV | 20 mV |
| $V_r$ – reset potential | 16 mV | 13 mV |
| $\tau$ – membrane time constant | 15 ms | 10 ms |
| $\tau_{ref}$ – absolute refractory period | 2 ms | 2ms |
| Synaptic STP parameters | Values |  |
| $U$ – probability of calcium binding | 0.2 | |
| $\tau_F$ – time constant of calcium unbinding | 1500 ms | |
| $\tau_D$ – time constant of transmitter refill | 200 ms | |
| Synaptic PRG1 parameters | Values |  |
| $M$ – probability of LPA binding | 0.6 | |
| $\tau_L$ – time constant of LPA unbinding | 30 s | |
| $\Delta U$ – change in $U$ during LPA binding | 0.1 | |
| Baseline input parameters | E | I |
| $\mu$ – mean external current | 23.1 mV/s | 21 mV/s |
| $\sigma$ – standard deviation of external current | 1 mV/s | 1 mV/s |
| Working memory input parameters | Values |  |
| $T_{init}$ – duration before cue stimulation | 10 s | |
| $T_{cue}$ – duration of cue stimulation | 350 ms | |
| $T_{total}$ – total duration of the simulation | 95 s | |
| $A_{cue}$ – contrast factor of selective cue | 1.15 | |

### Firing rate model

Analogously to the spiking network model, Table 3 depicts the firing rate network (see Figure 3). Implementation of the rate model and analysis of the data are performed with Matlab 2020a (code available upon request).

**Table 3:** Firing rate network model.

| Model summary |  |  |  |
| --- | --- | --- | --- |
| **Populations** | One: excitatory selective population $E_s$ (mean-field representation) | | |
| **Connectivity** | $E_s$ is connected to itself via an STP synapse with connection strength $J$ | | |
| **Neuron model** | — |  |  |
| **Synapse model** | Mean-field STP synapse (with or without noise) |  |  |
| **Input** | Constant (negative) input $E_{base}$ into population E | | |
| **Measurements** | Population activity $E$ of the network population $E_s$, state variables of the mean-field STP synapse | | |

| Populations |  |  |  |
| --- | --- | --- | --- |
| Name | Type | Dynamics | Function |
| $E_s$ | Mean-field representation of an excitatory neural population, characterized by its population activity $E$ | Population activity dynamics:<br>$\tau \frac{dE}{dt}(t) = -E + g(JyxE + E_{base}),$ where $\tau$ is the time constant of the population activity dynamics and $y$ and $x$ are the synaptic STP parameters for presynaptic calcium binding and neurotransmitter availability respectively. The nonlinear gain function $g$ is given by<br>$g(z) = \alpha \log(1 + \exp(z/\alpha)).$ | Excitatory population that is selective to the WM cue |

| Synapse models |  |
| --- | --- |
| Name | Dynamics |
| Noise-free STP synapse | Noise-free synaptic dynamics of the presynaptic calcium binding variable $y$ and the neurotransmitter availability $x$:<br>$\frac{dy}{dt} = \frac{1}{\tau_F}(U - y + U(1 - y)E)$ $\frac{dx}{dt} = \frac{1}{\tau_D}(1 - x - yxE),$<br>with parameters $U$ (rate of presynaptic calcium binding), $\tau_F$ (time constant of calcium unbinding) and $\tau_D$ (time constant of neurotransmitter replenishment). |
| Noisy STP synapse | Synaptic dynamics with noise:<br>$\frac{dy}{dt} = \frac{1}{\tau_F}(U - y + U(1 - y)E) + \xi(t)$ $\frac{dx}{dt} = \frac{1}{\tau_D}(1 - x - yxE),$<br>where $\xi(t)$ is a Gaussian white-noise perturbation with mean zero and autocorrelation $\langle \xi(t)\xi(t') \rangle = \sigma_y^2 \delta(t - t')$. |
| Input |  |
| Baseline input | Constant input $E_{\text{base}}$, takes negative values to reflect inputs from inhibitory populations not included in the model |
| Cue stimulation | After an initial period of baseline input stimulation $T_{\text{init}}$, the population $E$ receives an elevated input current $E_{\text{cue}} = A_{\text{cue}} \times E_{\text{base}}$ for a short period $T_{\text{cue}}$, which represents a stimulation of the network with a working memory cue. Afterwards, the external input is reduced to baseline level $E_{\text{base}}$ for the remainder of the simulation. |
| Measurements |  |
| Type | Method |
| Population activity $E$ | |
| Duration of population spike regime | Analogous to duration measurement in the spiking network |
| Working memory regime | Classification of delay activity as one of four dynamical regimes:<br>• silent – no population spikes during the delay, i.e. no synchronized spike with amplitude larger than 60 percent of the cue response<br>• transient – at least one population spike during delay and time between last observed spike and the end of the simulation larger than the maximum interspike interval (ISI)<br>• persistent – regular population spiking until the end of the simulation<br>• self-evoked – population spikes already before the cue presentation or extended periods without population spikes during the delay |

**Table 4:** Standard parameters of the firing rate network.

| **General simulation parameters** | **Values** |
| --- | --- |
| $dt$ – simulation time step | 0.1 ms |
| number of simulated realizations in Figure 3 D | 30 |
| **Population parameters** | **Values** |
| $\tau$ – time constant of population activity in population E | 13 ms |
| $J$ – synaptic efficacy of E-E synapse | 10 |
| $\alpha$ – exponent of nonlinear gain function $g$ | 1.5 |
| **Synaptic parameters** | **Values** |
| $U$ – probability of calcium binding | 0.2 |
| $\tau_F$ – time constant of calcium unbinding | 1500 ms |
| $\tau_D$ – time constant of transmitter refill | 200 ms |
| $\sigma_y$ – strength of white-noise perturbation on calcium binding variable $y$ in noisy STP synapses | 0 .. 0.005 |
| **Baseline input parameters** | **E** |
| $E_{\text{base}}$ – mean external current | -3.5 .. -2.5 |
| **Working memory input parameters** | **Values** |
| $T_{\text{init}}$ – duration before cue stimulation | 5 s |
| $T_{\text{cue}}$ – duration of cue stimulation | 350 ms |
| $T_{\text{total}}$ – total duration of the simulation | 29 s |
| $A_{\text{cue}}$ – contrast factor of selective cue | 1.15 |

## Supplementary Information

**Figure S1:**
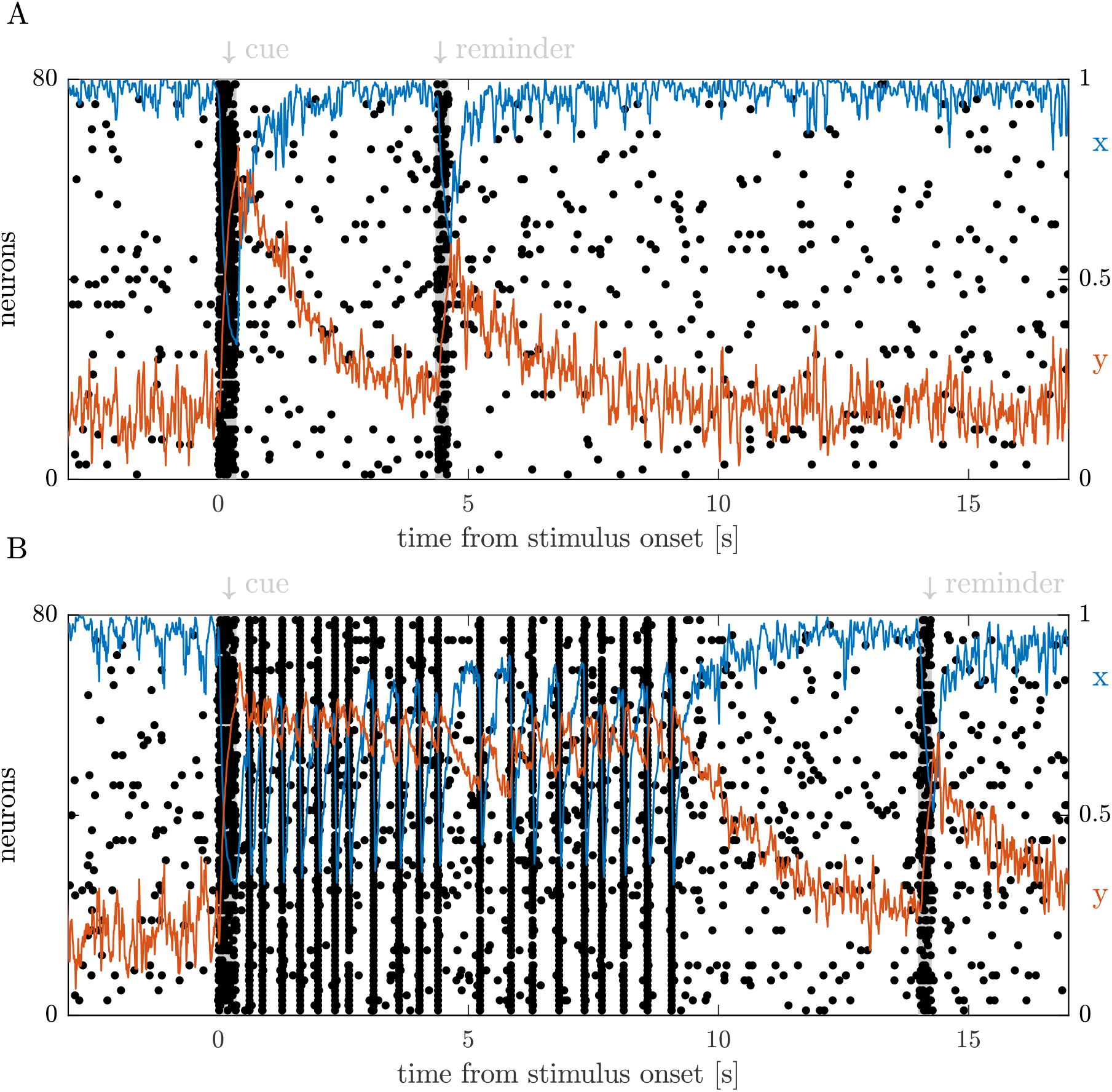
A weak reminder stimulus can re-evoke a population spike in the silent regime (A) and after transient WM activity (B) in spiking networks. After the end of the cue stimulation (A) and the transient WM activity (B), the synaptic neurotransmitter availability *x* and the presynaptic calcium binding *y* of spiking STP and PRG1 networks slowly decay back to their steady-state values. During the decay of these synaptic traces, a weak reminder stimulus into the excitatory selective population re-evokes a single population spike. Parameters: (A) standard PRG1 network (see Methods) with baseline input *E*_base_ = 22.95 mV/s and *τ*_*L*_ = 5 s as in Figure 2 A (left); (B) standard PRG1 network with baseline input *E*_base_ = 23.175 mV/s and *τ*_*L*_ = 5 s as in Figure 2 A (second from the left). A weak reminder stimulus (*E*_reminder_ = 1.05 *× E*_base_) is presented 4 s after the termination of cue stimulation (A) and 5 s after the end of transient activity (B) for 250 ms, respectively.

**Figure S2:**
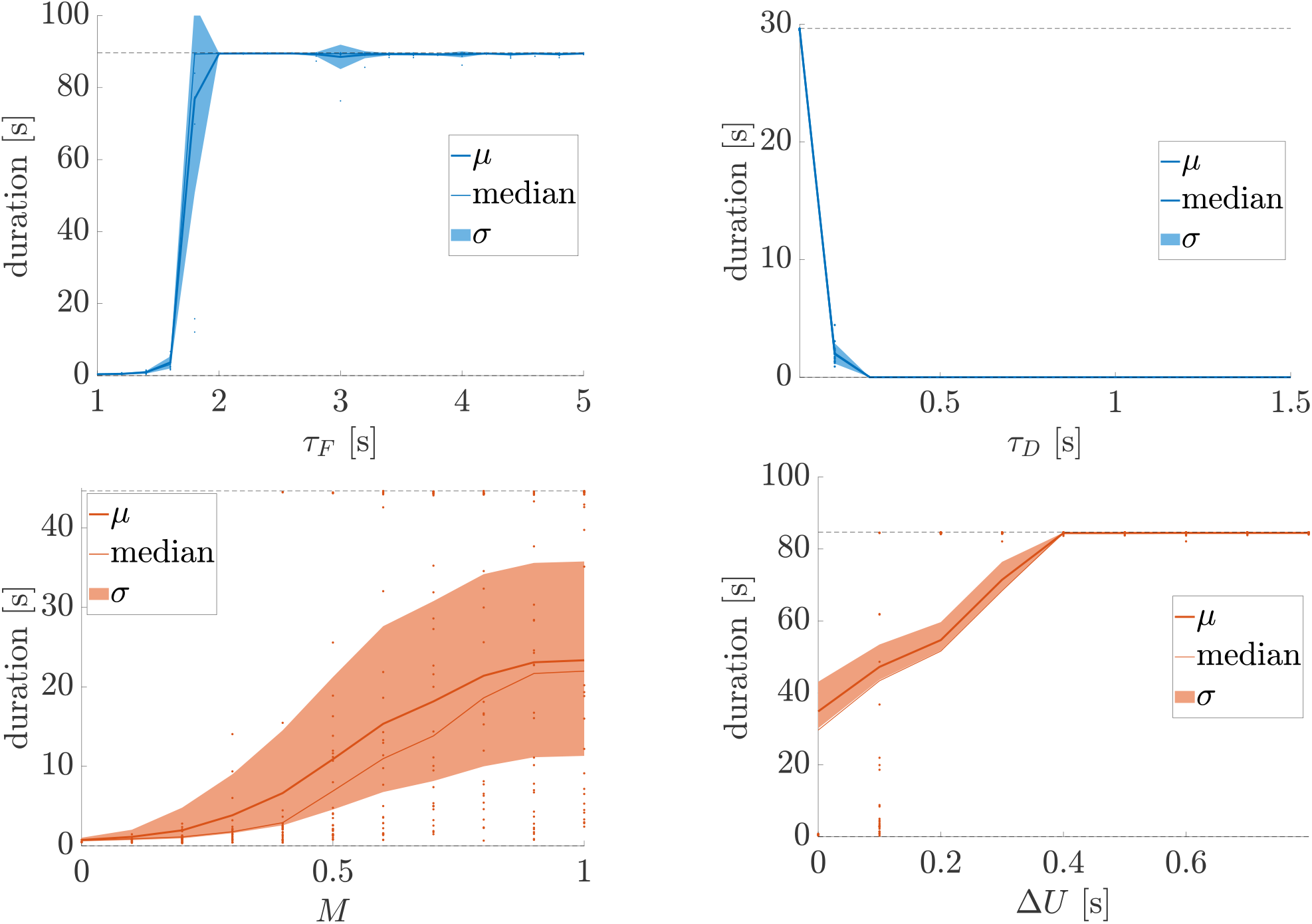
Modulation of transient WM duration by additional synaptic STP and PRG1 parameters. Transient WM durations are regulated by the synaptic STP parameters *τ*_*F*_ (top left, time constant of calcium unbinding) and *τ*_*D*_ (top right, time constant of neurotransmitter replenishment) and by the PRG1 parameters *M* (bottom left, LPA binding rate) and Δ*U* (bottom right, LPA-mediated change of the presynaptic calcium binding rate *U*). In Figure 2 B-D of the main manuscript, we show that the duration of transient WM activity is also controlled by the calcium binding rate *U* in STP networks and the time constant of LPA unbinding *τ*_*L*_ in PRG1 networks.

**Figure S3:**
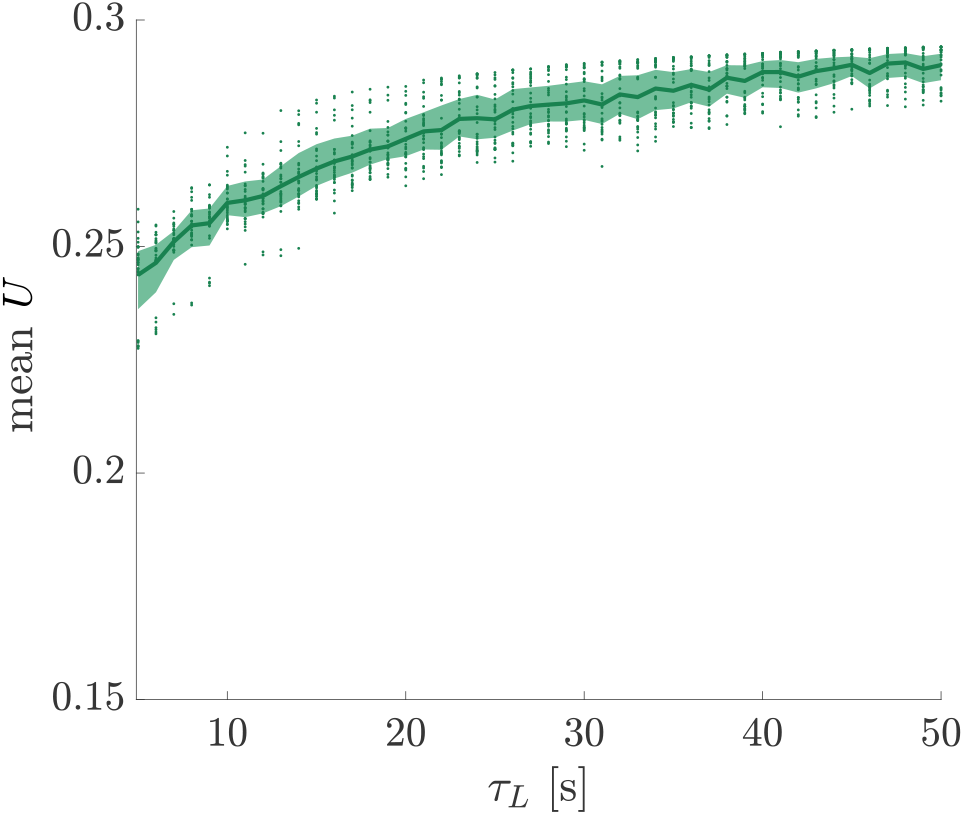
Time constant of LPA unbinding *τ*_*L*_ modulates transient WM duration via the presynaptic calcium binding rate *U*. For each value of the LPA-unbinding time constant *τ*_*L*_, we simulate 30 instances of a PRG1 network and compute the temporal average of the effective presynaptic calcium binding rate *U* = *U*_**base**_ + *l×* Δ*U* across the period of transient population spiking during the working memory delay. The average of *U* during the transient working memory regime increases with the synaptic timescale *τ*_*L*_ of LPA unbinding. Since *U* directly regulates the duration of the transient regime in STP networks (Figure 2 B), this suggests that *τ*_*L*_ controls transient durations in the PRG1 network (Figure 2 C, D) via its influence on *U*.

## References

Allen, N. J. and Eroglu, C. (2017), Cell Biology of Astrocyte-Synapse Interactions, Neuron 96(3), 697–708.

Baddeley, A. (2012), Working Memory: Theories, Models, and Controversies, Annu. Rev. Psychol. 63, 1–29.

Barbosa, J., Stein, H., Martinez, R. L., Galan-Gadea, A., Li, S., Dalmau, J., Adam, K. C., Valls-Solé, J., Constantinidis, C. and Compte, A. (2020), Interplay between persistent activity and activity-silent dynamics in the prefrontal cortex underlies serial biases in working memory, Nat. Neurosci. 23, 1016–1024.

Bastos, A. M., Loonis, R., Kornblith, S., Lundqvist, M. and Miller, E. K. (2018), Laminar recordings in frontal cortex suggest distinct layers for maintenance and control of working memory, Proc. Natl. Acad. Sci. U. S. A. 115(5), 1117–1122.

Belforte, J. E., Zsiros, V., Sklar, E. R., Jiang, Z., Yu, G., Li, Y., Quinlan, E. M. and Nakazawa, K. (2010), Postnatal NMDA receptor ablation in corticolimbic interneurons confers schizophrenia-like phenotypes, Nat. Neurosci. 13(1), 76–83.

Berridge, M. J., Bootman, M. D. and Roderick, H. L. (2003), Calcium signalling: dynamics, homeostasis and remodelling, Nat. Rev. Mol. Cell Biol. 4(7), 517–529.

Bouchacourt, F. and Buschman, T. J. (2019), A Flexible Model of Working Memory, Neuron 103, 147–160.

Bräuer, A. U., Savaskan, N. E., Kühn, H., Prehn, S., Ninnemann, O. and Nitsch, R. (2003), A new phospholipid phosphatase, PRG-1, is involved in axon growth and regenerative sprouting, Nat. Neurosci. 6(6), 572–578.

Brunel, N. (2000), Dynamics of Sparsely Connected Networks of Excitatory and Inhibitory Spiking Neurons, J. Comput. Neurosci. 8, 183–208.

Buschman, T. J., Siegel, M., Roy, J. E. and Miller, E. K. (2011), Neural substrates of cognitive capacity limitations, Proc. Natl. Acad. Sci. 108(27), 11252–11255.

Christophel, T. B., Christiaan Klink, P., Spitzer, B., Roelfsema, P. R. and Haynes, J.-D. (2017), The Distributed Nature of Working Memory, Trends Cogn. Sci. 21(2), 111–124.

Christopher, G. and MacDonald, J. (2005), The impact of clinical depression on working memory, Cogn. Neuropsychiatry 10(5), 379–399.

Cortes, J. M., Desroches, M., Rodrigues, S., Veltz, R., Muñoz, M. A. and Sejnowski, T. J. (2013), Short-term synaptic plasticity in the deterministic Tsodyks-Markram model leads to unpredictable network dynamics, Proc. Natl. Acad. Sci. U. S. A. 110(41), 16610–16615.

Coyle, J. T. (2006), Glutamate and schizophrenia: beyond the dopamine hypothesis., Cell. Mol. Neurobiol. 26(4-6), 365–384.

De Pittà, M., Brunel, N. and Volterra, A. (2016), Astrocytes: Orchestrating synaptic plasticity?, Neuroscience 323, 43–61.

Druckmann, S. and Chklovskii, D. B. (2012), Neuronal circuits underlying persistent representations despite time varying activity, Curr. Biol. 22, 2095–2103.

Edin, F., Klingberg, T., Johansson, P., McNab, F., Tegnér, J. and Compte, A. (2009), Mechanism for top-down control of working memory capacity, Proc. Natl. Acad. Sci. 106(16), 6802–6807.

Fiebig, F. and Lansner, A. (2017), A spiking working memory model based on hebbian short-term potentiation, J. Neurosci. 37(1), 83–96.

Fioravanti, M., Bianchi, V. and Cinti, M. E. (2012), Cognitive deficits in schizophrenia: An updated metanalysis of the scientific evidence, BMC Psychiatry 12(64).

Freeman, W. J., Chang, H. J., Burke, B. C., Rose, P. A. and Badler, J. (1997), Taming chaos: Stabilization of aperiodic attractors by noise, IEEE Trans. Circuits Syst. I Fundam. Theory Appl. 44(10), 989–996.

Frydecka, D., Eissa, A. M., Hewedi, D. H., Ali, M., DrapaÅ‚a, J., Misiak, B., KÅ‚osiÅ„ska, E., Phillips, J. R. and Moustafa, A. A. (2014), Impairments of working memory in schizophrenia and bipolar disorder: the effect of history of psychotic symptoms and different aspects of cognitive task demands, Front. Behav. Neurosci. 8, 416.

Fujisawa, S., Amarasingham, A., Harrison, M. T. and Buzsáki, G. (2008), Behavior-dependent short-term assembly dynamics in the medial prefrontal cortex, Nat. Neurosci. 11(7), 823 – 833.

Funahashi, S., Bruce, C. J. and Goldman-Rakic, P. S. (1989), Mnemonic Coding of Visual Space in the Monkey’s Dorsolateral Prefrontal Cortex, J. Neurophysiol. 61(2), 331–349.

Fuster, J. M. and Alexander, G. E. (1971), Neuron Activity Related to Short-Term Memory, Science 173(3997), 652–654.

Gravitz, L. (2019), The Importance of Forgetting, Nature 571, S12–S14.

Harrison, P. J. and Weinberger, D. R. (2005), Schizophrenia genes, gene expression, and neuropathology: On the matter of their convergence, Mol. Psychiatry 10(1), 40–68.

Harvey, C. D., Coen, P. and Tank, D. W. (2012), Choice-specific sequences in parietal cortex during a virtual-navigation decision task, Nature 484(7392), 62–68.

Hu, Y., Brunton, S. L., Cain, N., Mihalas, S., Kutz, J. N. and Shea-Brown, E. (2018), Feed-back through graph motifs relates structure and function in complex networks, Phys. Rev. E 98(6), 1–25.

Jonides, J., Lewis, R. L., Nee, D. E., Lustig, C. A., Berman, M. G. and Moore, K. S. (2008), The Mind and Brain of Short-Term Memory, Annu. Rev. Psychol. 59(1), 193–224.

Juel, A., Darbyshire, A. G. and Mullin, T. (1997), The effect of noise on pitchfork and Hopf bifurcations, Proc. R. Soc. London A 453, 2627–2647.

Kasper, L. J., Alderson, R. M. and Hudec, K. L. (2012), Moderators of working memory deficits in children with attention-deficit/hyperactivity disorder (ADHD): A meta-analytic review, Clin. Psychol. Rev. 32(7), 605–617.

Kercood, S., Grskovic, J. A., Banda, D. and Begeske, J. (2014), Working memory and autism: A review of literature, Res. Autism Spectr. Disord. 8(10), 1316–1332.

Keune, W.-J., Hausmann, J., Bolier, R., Tolenaars, D., Kremer, A., Heidebrecht, T., Joosten, R. P., Sunkara, M., Morris, A. J., Matas-Rico, E., Moolenaar, W. H. et al. (2016), Steroid binding to Autotaxin links bile salts and lysophosphatidic acid signalling, Nat. Commun. 7(1), 11248.

Kiss, I. Z., Hudson, J. L., Escalera Santos, G. J. and Parmananda, P. (2003), Experiments on coherence resonance: Noisy precursors to Hopf bifurcations, Phys. Rev. E 67(3), 035201.

Kolata, S., Wu, J., Light, K., Schachner, M. and Matzel, L. D. (2008), Impaired working memory duration but normal learning abilities found in mice that are conditionally deficient in the close homolog of L1, J. Neurosci. 28(50), 13505–13510.

LaRocque, J. J., Lewis-Peacock, J. A. and Postle, B. R. (2014), Multiple neural states of representation in short-term memory? It’s a matter of attention, Front. Hum. Neurosci. 8, 5.

Lee, J. and Lee, J. (2018), Quantitative analysis of a transient dynamics of a gene regulatory network, Phys. Rev. E 98(6), 062404.

Lewis, D. A., Volk, D. W. and Hashimoto, T. (2004), Selective alterations in prefrontal cortical GABA neurotransmission in schizophrenia: A novel target for the treatment of working memory dysfunction, Psychopharmacology 174(1), 143–150.

Lovinger, D. M. (2008), Presynaptic Modulation by Endocannabinoids, in T. Südhof and K. Starke, eds, ‘Pharmacol. Neurotransmitter Release. Handb. Exp. Pharmacol.’, Springer, Berlin, Heidelberg, Berlin, Heidelberg, chapter vol. 184, pp. 435–477.

Lundqvist, M., Herman, P., Warden, M. R., Brincat, S. L. and Miller, E. K. (2018), Gamma and beta bursts during working memory readout suggest roles in its volitional control, Nat. Commun. 9(1), 1–12.

Lundqvist, M., Rose, J., Herman, P., Brincat, S. L. L., Buschman, T. J. J. and Miller, E. K. K. (2016), Gamma and Beta Bursts Underlie Working Memory, Neuron 90(1), 152–164.

Markram, H., Muller, E., Ramaswamy, S., Reimann, M. W., Abdellah, M., Sanchez, C. A., Ailamaki, A., Alonso-Nanclares, L., Antille, N., Arsever, S. et al. (2015), Reconstruction and Simulation of Neocortical Microcircuitry, Cell 163(2), 456–492.

Masse, N. Y., Yang, G. R., Francis Song, H., Wang, X.-J. and Freedman, D. J. (2019), Circuit mechanisms for the maintenance and manipulation of information in working memory, Nat. Neurosci. 22, 1159–1167.

Mastrogiuseppe, F. and Ostojic, S. (2018), Linking Connectivity, Dynamics, and Computations in Low-Rank Recurrent Neural Networks, Neuron 99(3), 609–623.e29.

Matt Alderson, R., Kasper, L. J., Hudec, K. L. and Patros, C. H. (2013), Attention-deficit/hyperactivity disorder (ADHD) and working memory in adults: A meta-analytic review, Neuropsychology 27(3), 287–302.

Mi, Y., Katkov, M. and Tsodyks, M. (2017), Synaptic Correlates of Working Memory Capacity, Neuron 93(2), 323–330.

Mongillo, G., Barak, O. and Tsodyks, M. (2008), Synaptic Theory of Working Memory, Science 319(5869), 1543–1546.

Moresco, E. M. Y., Scheetz, A. J., Bornmann, W. G., Koleske, A. J. and Fitzsimonds, R. M. (2003), Abl Family Nonreceptor Tyrosine Kinases Modulate Short-Term Synaptic Plasticity, J. Neurophysiol. 89(3), 1678–1687.

Morrone, J. A., Markland, T. E., Ceriotti, M. and Berne, B. J. (2011), Efficient multiple time scale molecular dynamics: Using colored noise thermostats to stabilize resonances, J. Chem. Phys. 134(1), 014103.

Murray, J. D., Jaramillo, J. and Wang, X.-J. (2017), Working Memory and Decision-Making in a Frontoparietal Circuit Model, J. Neurosci. 37(50), 12167–12186.

Nordlie, E., Gewaltig, M. O. and Plesser, H. E. (2009), Towards reproducible descriptions of neuronal network models, PLoS Comput. Biol. 5(8), e1000456.

Olivers, C. N., Peters, J., Houtkamp, R. and Roelfsema, P. R. (2011), Different states in visual working memory: When it guides attention and when it does not, Trends Cogn. Sci. 15(7), 327–334.

Orhan, A. E. and Ma, W. J. (2019), A diverse range of factors affect the nature of neural representations underlying short-term memory, Nat. Neurosci. 22, 275–283.

Pasternak, T. and Greenlee, M. W. (2005), Working memory in primate sensory systems, Nat. Rev. Neurosci. 6, 97–107.

Postle, B. R. (2006), Working memory as an emergent property of the mind and brain, Neuroscience 139(1), 23–38.

Rose, E. J. and Ebmeier, K. P. (2006), Pattern of impaired working memory during major depression, J. Affect. Disord. 90(2-3), 149–161.

Schmitt, L. I., Wimmer, R. D., Nakajima, M., Happ, M., Mofakham, S. and Halassa, M. M. (2017), Thalamic amplification of cortical connectivity sustains attentional control, Nature 545(7653), 219–223.

Schuessler, F., Dubreuil, A., Mastrogiuseppe, F., Ostojic, S. and Barak, O. (2020), Dynamics of random recurrent networks with correlated low-rank structure, Phys. Rev. Res. 2(1), 13111.

Shilyansky, C., Karlsgodt, K. H., Cummings, D. M., Sidiropoulou, K., Hardt, M., James, A. S., Ehninger, D., Bearden, C. E., Poirazi, P., Jentsch, J. D. et al. (2010), Neurofibromin regulates corticostriatal inhibitory networks during working memory performance, Proc. Natl. Acad. Sci. 107(29), 13141–13146.

Stokes, M. G. (2015), ’Activity-silent’ working memory in prefrontal cortex: A dynamic coding framework, Trends Cogn. Sci. 19(7), 394–405.

Sun, X.-D., Li, L., Liu, F., Huang, Z.-H., Bean, J. C., Jiao, H.-F., Barik, A., Kim, S.-M., Wu, H., Shen, C. et al. (2016), Lrp4 in astrocytes modulates glutamatergic transmission, Nat. Neurosci. 19(8), 1010–1018.

Thalman, C., Horta, G., Qiao, L., Endle, H., Tegeder, I., Cheng, H., Laube, G., Sigrudsson, T., Hauser, M. J., Tenzer, S. et al. (2018), Synaptic phospholipids as a new target for cortical hyperexcitability and E/I balance in psychiatric disorders, Nat. Mol. Psychiatry 23(8), 1699–1710.

Tokumitsu, H., Hatano, N., Tsuchiya, M., Yurimoto, S., Fujimoto, T., Ohara, N., Kobayashi, R. and Sakagami, H. (2010), Identification and characterization of PRG-1 as a neuronal calmodulin-binding protein, Biochem. J. 431(1), 81–91.

Trimbuch, T., Beed, P., Vogt, J., Schuchmann, S., Maier, N., Kintscher, M., Breustedt, J., Schuelke, M., Streu, N., Kieselmann, O. et al. (2009), Synaptic PRG-1 Modulates Excitatory Transmission via Lipid Phosphate-Mediated Signaling, Cell 138(6), 1222–1235.

Trübutschek, D., Marti, S., Ojeda, A., King, J. R., Mi, Y., Tsodyks, M. and Dehaene, S. (2017), A theory of working memory without consciousness or sustained activity, Elife 6, e23871.

Trübutschek, D., Marti, S., Ueberschär, H. and Dehaene, S. (2019), Probing the limits of activity-silent non-conscious working memory, Proc. Natl. Acad. Sci. U. S. A. 116(28), 14358–14367.

Tsodyks, M., Pawelzik, K. and Markram, H. (1998), Neural Networks with Dynamic Synapses, Neural Comput. 10(4), 821–835.

Unichenko, P., Kirischuk, S., Yang, J. W., Baumgart, J., Roskoden, T., Schneider, P., Sommer, A., Horta, G., Radyushkin, K., Nitsch, R. et al. (2016), Plasticity-related gene 1 affects mouse barrel cortex function via strengthening of glutamatergic thalamocortical transmission, Cereb. Cortex 26(7), 3260–3272.

Ushakov, O. V., Wünsche, H.-J., Henneberger, F., Khovanov, I. A., Schimansky-Geier, L. and Zaks, M. A. (2005), Coherence resonance near a Hopf bifurcation, Phys. Rev. Lett. 95(12), 123903.

Van Snellenberg, J. X., Girgis, R. R., Horga, G., van de Giessen, E., Slifstein, M., Ojeil, N., Weinstein, J. J., Moore, H., Lieberman, J. A., Shohamy, D. et al. (2016), Mechanisms of Working Memory Impairment in Schizophrenia, Biol. Psychiatry 80(8), 617–626.

Van Vreeswijk, C. and Sompolinsky, H. (1998), Chaotic Balanced State in a Model of Cortical Circuits, Neural Comput. 10(6), 1321–1371.

Vogt, J., Kirischuk, S., Unichenko, P., Schlüter, L., Pelosi, A., Endle, H., Yang, J.-W., Schmarowski, N., Cheng, J., Thalman, C. et al. (2017), Synaptic Phospholipid Signaling Modulates Axon Outgrowth via Glutamate-dependent Ca2+-mediated Molecular Pathways, Cereb. Cortex 27(1), 131–145.

Vogt, J., Yang, J.-W., Mobascher, A., Cheng, J., Li, Y., Liu, X., Baumgart, J., Thalman, C., Kirischuk, S., Unichenko, P. et al. (2016), Molecular cause and functional impact of altered synaptic lipid signaling due to a prg-1 gene SNP, EMBO Mol. Med. 8(1), 25–38.

Von Engelhardt, J., Mack, V., Sprengel, R., Kavenstock, N., Li, K. W., Stern-Bach, Y., Smit, A. B., Seeburg, P. H. and Monyer, H. (2010), CKAMP44: A brain-specific protein attenuating short-term synaptic plasticity in the dentate gyrus, Science 327(5972), 1518–1522.

Watanabe, K. and Funahashi, S. (2014), Neural mechanisms of dual-task interference and cognitive capacity limitation in the prefrontal cortex, Nat. Neurosci. 17(4), 601–611.

Wei, Y. and Koulakov, A. A. (2014), Long-term memory stabilized by noise-induced rehearsal, J. Neurosci. 34(47), 15804–15815.

Yizhar, O., Fenno, L. E., Prigge, M., Schneider, F., Davidson, T. J., Ogshea, D. J., Sohal, V. S., Goshen, I., Finkelstein, J., Paz, J. T. et al. (2011), Neocortical excitation/inhibition balance in information processing and social dysfunction, Nature 477(7363), 171–178.

Yung, Y. C., Stoddard, N. C., Mirendil, H. and Chun, J. (2015), Lysophosphatidic Acid Signaling in the Nervous System, Neuron 85(4), 669–682.

Zucker, R. S. and Regehr, W. G. (2002), Short-Term Synaptic Plasticity, Annu. Rev. Physiol. 64(1), 355–405.

